# Allelic and Genotypic Frequencies of NAT2, CYP2E1 and AADAC genes in a cohort of Peruvian Tuberculosis Patients

**DOI:** 10.1101/2021.02.27.433209

**Authors:** Kelly S. Levano, Luis Jaramillo-Valverde, David D. Tarazona, Cesar Sanchez, Silvia Capristano, Lely Solari, Alberto Mendoza-Ticona, Alonso Soto, Christian Rojas, Roberto Zegarra-Chapoñan, Heinner Guio

## Abstract

**Background:** We determined the frequency of genetic polymorphisms in three anti-TB drug metabolic proteins previously reported: N-acetyltransferase 2 (NAT2), cytochrome P450 2E1 (CYP2E1) and arylacetamide deacetylase (AADAC) within a Peruvian population in a cohort of TB patients. We included 395 participants completed their anti-tuberculosis treatment.

**Results:** ∼74% of the participants are carriers of slow metabolizer genotypes: NAT2*5, NAT2*6 and NAT2*7, which increase the sensitivity of INH at low doses and increase the risk of drug-induced liver injuries. ∼ 64% are homozygous for the wild-type CYP2E1*1A allele, which could increase the risk of hepatotoxicity. However, 16% had a NAT2 fast metabolizer phenotype which could increase the risk of acquiring resistance to INH, thereby increasing the risk of multidrug-resistant (MDR) or treatment failure. The frequency of rs1803155 (AADAC*2 allele) was higher (99.9%) in Peruvians than in in European American, African American, Japanese, and Korean populations.

**Conclusions:** This high prevalence of slow metabolizers for Isoniazid in the Peruvian population should be further studied and considered to help individualize drug regimens, especially in countries with a great genetic diversity like Peru. These data will help the Peruvian National Tuberculosis Control Program develop new strategies for therapies.

## Background

Tuberculosis (TB) continues to be a leading cause of global morbidity and mortality, with about 10 million cases and a total of 1.2 million deaths reported in 2019 [1]. Even though the current TB regimen is highly effective under optimal conditions, there are still many undefined issues including drug underexposure, high prevalence if drug-related toxicity, selection of resistant strains and variability of response [2], which could be explained by the variability in the pharmacokinetics of anti-TB drugs. Mutations or polymorphisms in genes encoding metabolic enzymes, transporters or carries can lead to this variability in drug pharmacokinetics and pharmacodynamics. The identification of these genetic variations could help select the right anti-TB drug, with the right dosage increasing efficacy and reducing drug-related toxicity and preventing drug resistance [2, 3]. To determine if genetic variabilities affecting drug response were present in the Peruvian population, which has a high TB burden with an estimated 32,970cases in 2019 [1], including a high prevalence of drug-resistant TB cases, we determined the frequency of genetic polymorphisms in three anti-TB drug metabolic proteins previously reported: N-acetyltransferase 2 (NAT2), cytochrome P450 2E1 (CYP2E1) and arylacetamide deacetylase (AADAC) [4]. These three proteins participate in the metabolism of the initial phase anti-TB drugs: isoniazid and rifampicin. In the liver, isoniazid is acetylated to its major metabolite, N-acetyl-isoniazid by the action of NAT2. It is then further deactivated by other enzymes including CYP2E1 [5, 6]. Thus, genetic variations in these two enzymes, leading to alterations in their enzymatic functions can cause variations in isoniazid pharmacokinetics. AADAC is one of the few known enzymes responsible for the deacetylation of rifampicin and AADAC allele decreased enzyme activity [7, 8]. Thus, genetic variations in these three enzymes, leading to alterations in their enzymatic functions could cause variations in isoniazid and rifampicin pharmacokinetics. For the present study, we selected previously reported single nucleotide polymorphisms (SNPs) that could alter NAT2, CYP2E1 and AADAC enzyme activity and determine their frequency within a Peruvian population in a cohort of TB patients.

## Methods

### Studied populations

Our study includes 395 unrelated individuals diagnosed with pulmonary tuberculosis between 2014-2015 recruited from health establishments of the Minister of Health (MINSA) located in Lima and Callao, Peru. The 395 participants (217 males and 178 females) completed their anti-tuberculosis treatment. Our study was approved by the Ethics in Research Committee of the Peruvian National Institute of Health (INS), and written informed consent was obtained from all the participants.

### Genotyping of NAT2, CYP2E1 and AADAC

Genomic DNA was extracted from peripheral blood of all 395 participants using the genomic DNA extraction kit QIAamp DNA Blood Mini Kit (Qiagen, Germany). The selected genomic DNA regions for the analysis of each gene included the most common reported SNPs (For NAT2: rs1041983, rs1801280, rs1799929, rs1799930, rs1208 and rs1799931; for CYP2E1: rs3813867 and rs2031920; for AADAC: rs1803155). These regions were amplified by PCR using Platinum Taq DNA polymerase kit (Invitrogen, USA) using the following primers: For NAT2: 5’-GTCACACGAGGAAATCAAATGCT-3 and 5’-CGTGAGGGTAGAGAGGATATCTG-3’; for CYP2E1: 5’-CCGTGAGCCAGTCGAGTCTA-3’ and 5’-TTCATTCTGTCTTCTAACTGGCAA-3’; and for AADAC: 5’-TCATTCCTAGCAGAAAGGAGATT-3’ and 5’-GCTCACATTTATTCTCTTGCATCG-3’. PCR-amplified fragments were purified using QIAmp Gel Purification Kit (Qiagen, USA). SNP genotyping on the purified fragments was performed using sanger sequencing (Macrogen, South Korea). Nucleotide substitutions were identified and analyzed using the Geneious version 9.1.5 (Biomatters Ltd., New Zealand).

### computational phenotyping for NAT2

Predicted phenotypes were determined from genotypes as three types of metabolizers: slow metabolizer (two slow alleles), rapid metabolizer (two rapid alleles) and intermediate metabolizer (one slow and another rapid acetylator allele). The alleles considered rapid were: wild-type NAT2∗4, 282C>T (NAT2∗13), 481C>T (NAT2∗11) and 803A>G (NAT2∗12), while the alleles considered slow were: 341T>C (NAT2∗5), 590G>A (NAT2∗6), 857G>A (NAT2∗7), and 191G>A (NAT2∗14) [9]. The computational inferred phenotypes using a combination of NAT2 SNPs for the 395 participants were determined using an online software program, NAT2PRED (nat2pred.rit.albany.edu) [10, 11].

### Statistical analysis

Genotype frequencies in our population were calculated in accordance with the Hardy-Weinberg equilibrium. Intervals confidence 95% were calculated for phenotypic genotypic and allelic frequencies. Data analysis was carried out using Stata 15 program (StataCorp. 2016. Stata Statistical Software: Release 15. College Station, TX, USA).

## Results

In this study, we determined the presence of the six most common SNPs, rs1041983 (282C>T), rs1801280 (341T>C), rs1799929 (481C>T), rs1799930 (590G>A), rs1208 (803A>G) and rs1799931(857G>A), of the NAT2 gene in the 395 individuals from Lima and Callao, Peru. No new SNPs were identified, indicating that the NAT2 gene has no other SNPs in the Peruvian population studied. The allele frequencies of these major NAT2 SNPs are represented in Table 1. This study found that NAT2∗13, C283T, is the most frequent genetic variant (∼40% of alleles) among our samples. The allele not harboring any mutation (wild-type NAT2∗4) was present in 44 of the 395 samples (∼11% of alleles). Table 2 shows the frequencies of NAT2 genotype obtained from the studied population. The most frequent observed heterozygote was also NAT2∗13 C283T (48.6%) followed by NAT2∗11 C481T (34.9%) and NAT2∗5 T341C (34.7%) among the Peruvian population studied. The lowest frequency of observed heterozygote genotypes was NAT2∗12 A803G with a frequency of 18.7%. In homozygote, the NAT2∗12 803A>G (31.2%) genotype was the most common one but the lowest homozygote among them was NAT2∗6 590G>A (12.5%). The linkage disequilibrium (LD) analysis is shown in Figure 1. The six NAT2 variants, 282C>T, 341T>C, 481C>T, 590G>A, 803A>G, and 857G>A were applied to Haploview software. The LD for each pair of genetic variants was measured using ∣D⍰ ∣ and correlation coefficient (r2 > 0:8). A haplotype **block** was found in the following SNP positons 341T>C and 481C>T (D :0,808 and r2:0,437) in Peruvian population samples which is identified as like strong LD. There were no significant differences observed between NAT2 genotypes with respect to age and gender. The NAT2 inferred metabolizing status was predicted using the 6 SNPs analyzed as stated above. As a result, the predicted metabolizing phenotype of fast, intermediate, and slow metabolizers was 14.9%, 38.2% and 46.8%, respectively (Table 3).

**Table 1:**
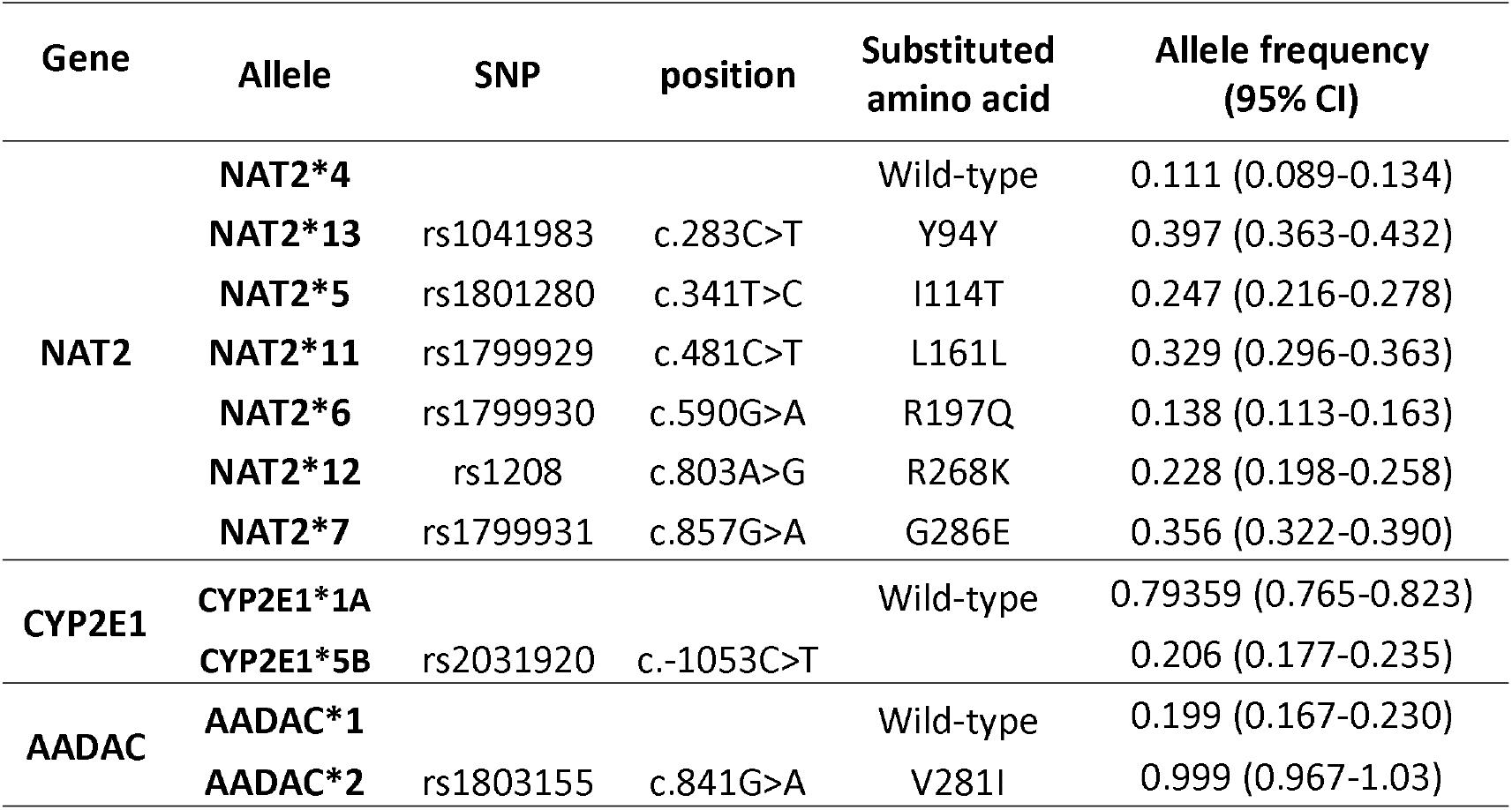
Allele frequencies of NAT2, CYP2E1 and AADAC polymorphisms in a Peruvian population (n = 395)

**Table 2:**
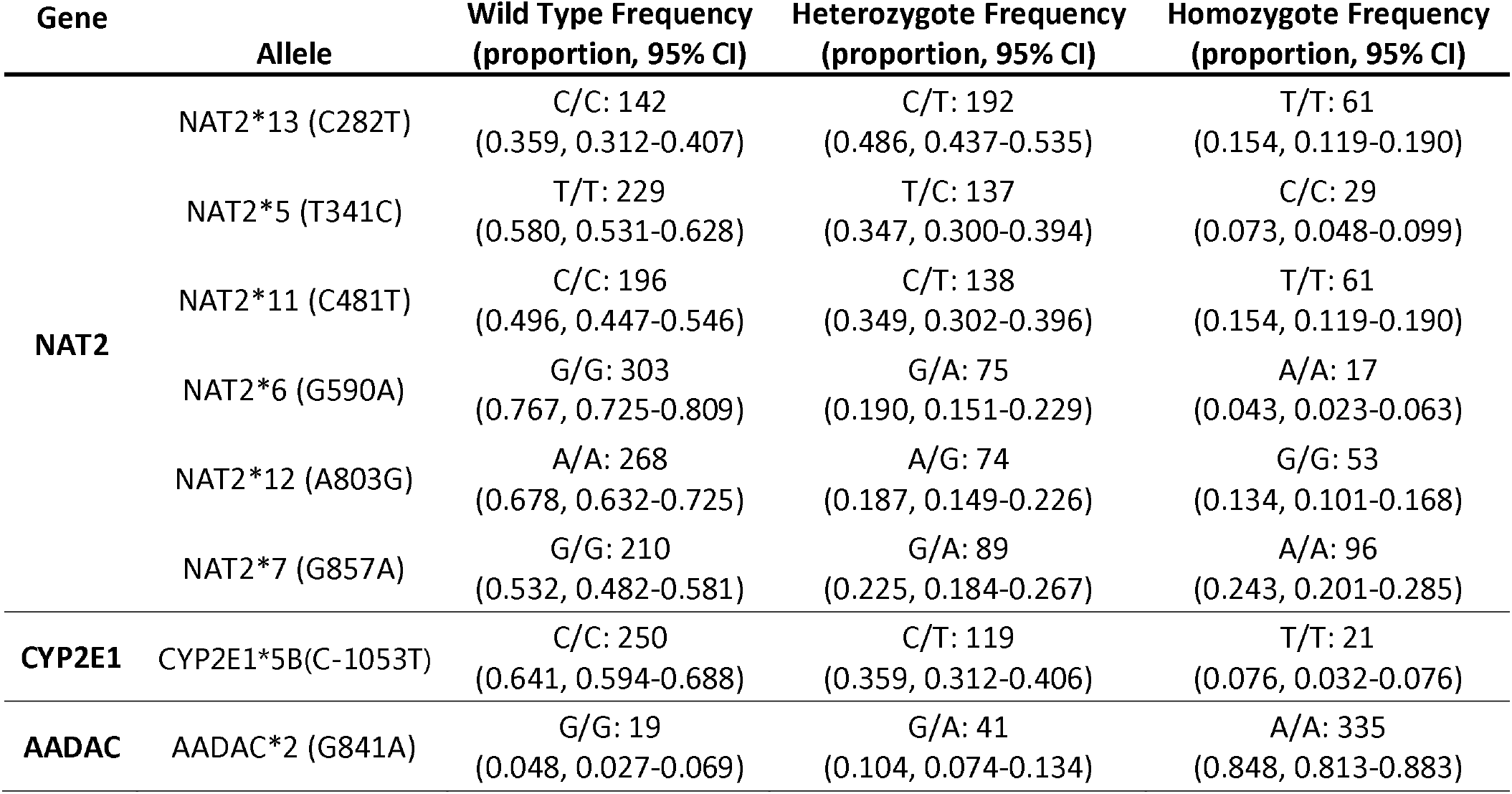
Genotype frequency of NAT2, CYP2E1 and AADAC genes in a Peruvian population (n = 395)

| Gene | Allele | Wild Type Frequency<br>(proportion, 95% CI) | Heterozygote Frequency<br>(proportion, 95% CI) | Homozygote Frequency<br>(proportion, 95% CI) |
| --- | --- | --- | --- | --- |
| NAT2 | NAT2*13 (C282T) | C/C: 142<br>(0.359, 0.312-0.407) | C/T: 192<br>(0.486, 0.437-0.535) | T/T: 61<br>(0.154, 0.119-0.190) |
|  | NAT2*5 (T341C) | T/T: 229<br>(0.580, 0.531-0.628) | T/C: 137<br>(0.347, 0.300-0.394) | C/C: 29<br>(0.073, 0.048-0.099) |
|  | NAT2*11 (C481T) | C/C: 196<br>(0.496, 0.447-0.546) | C/T: 138<br>(0.349, 0.302-0.396) | T/T: 61<br>(0.154, 0.119-0.190) |
|  | NAT2*6 (G590A) | G/G: 303<br>(0.767, 0.725-0.809) | G/A: 75<br>(0.190, 0.151-0.229) | A/A: 17<br>(0.043, 0.023-0.063) |
|  | NAT2*12 (A803G) | A/A: 268<br>(0.678, 0.632-0.725) | A/G: 74<br>(0.187, 0.149-0.226) | G/G: 53<br>(0.134, 0.101-0.168) |
|  | NAT2*7 (G857A) | G/G: 210<br>(0.532, 0.482-0.581) | G/A: 89<br>(0.225, 0.184-0.267) | A/A: 96<br>(0.243, 0.201-0.285) |
| CYP2E1 | CYP2E1*5B(C-1053T) | C/C: 250<br>(0.641, 0.594-0.688) | C/T: 119<br>(0.359, 0.312-0.406) | T/T: 21<br>(0.076, 0.032-0.076) |
| AADAC | AADAC*2 (G841A) | G/G: 19<br>(0.048, 0.027-0.069) | G/A: 41<br>(0.104, 0.074-0.134) | A/A: 335<br>(0.848, 0.813-0.883) |

**Table 3:** Predicted metabolizing phenotype for NAT2 in a Peruvian population (n = 395)

| Gene | Metabolizing Phenotype * |  |  |
| --- | --- | --- | --- |
|  | Proportion (95% CI) |  |  |
| NAT2 | Fast | Intermediate | Slow |
|  | 0.149 | 0.382 | 0.468 |
|  | (0.114-0.185) | (0.334-0.430) | (0.419-0.518) |

\*The metabolizing phenotype was determined using the online software
http://nat2pred.rit.albany.edu/.

**Table 4:** Distribution of NAT2 alleles among the Peruvian population studied compared with various human population.

| Population | Peru<br>(current<br>study) | Brazil<br>(2016) | Mexico<br>(2012) | Spain<br>(2011) |
| --- | --- | --- | --- | --- |
| NAT2*4 (Wild-type) | 0.111 | 0.258 | 0.306 | 0.186 |
| NAT2*13 (C282T) | 0.099 | 0.008 | 0.008 | - |
| NAT2*5 (T341C) | 0.420 | 0.446 | 0.312 | 0.417 |
| NAT2*11 (C481T) | 0.035 | - | - | - |
| NAT2*6 (G590A) | 0.099 | 0.150 | 0.174 | 0.292 |
| NAT2*12 (A803G) | 0.013 | 0.023 | 0.048 | - |
| NAT2*7 (G857A) | 0.223 | 0.096 | 0.140 | 0.106 |

**Figure 1.**
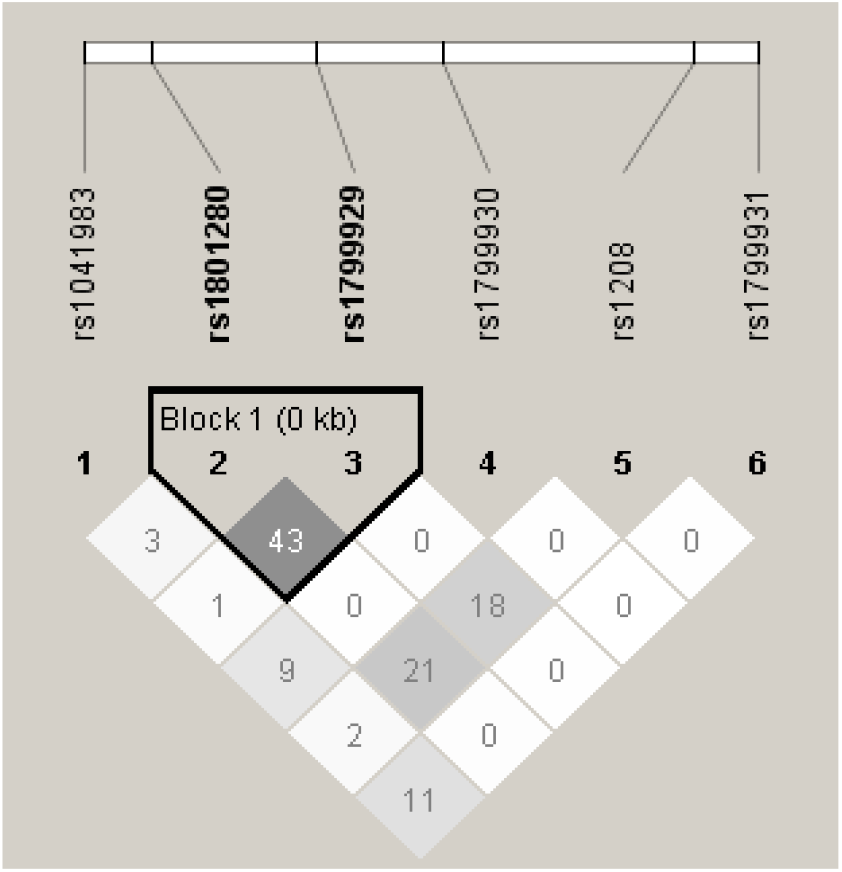
Linkage disequilibrium for NAT2 metabolizer-associated SNPs in Peruvian population studied.

The allelic and genotypic distribution of the rs2031920 (-1053C>T) variant of CYP2E1 among the studied population is shown in Tables 1 and 2, respectively. The results show an allele frequency of ∼21% for the CYP2E1 variant, and ∼79% for the wild-type allele. The observed genotype frequency of the homozygous and heterozygotes were ∼7% and ∼36%, respectively.

We also analyzed the allele and genotype frequency of the rs1803155 SNP of the AADAC gene in the 395 individuals of this study (Table 1 and 2). According to our results, the AADAC genetic variant has an allele frequency of ∼99.9%, while the wild-type allele was ∼20%. The homozygous and heterozygous genotype distribution were ∼85% and ∼10%, respectively.

## Discussion

Studies in different populations have shown ethnic variabilities in both NAT2 and CYP2E1 genotypes and phenotypes. There is still limited information about the genetic variations in the Peruvian population. In the current study, we analyzed NAT2, CYP2E1 and AADAC genotypes and allele frequencies in 395 individuals from Peru. As stated above, NAT2 and CYP2E1 are two essential enzymes in the metabolism of INH. Altered NAT2 and/or CYP2E1 activities due to polymorphic genotypes can result in (a) the accumulation of toxic substances in the liver, and (b) variations in INH plasma concentrations that can affect the efficacy of the drug.

In the analysis of the *NAT2* gene, the results showed that *NAT2*13* (39.7% of alleles) and *NAT2*7* (35.6% of alleles) were the most frequent genetic variants amount the population studied. *NAT2*13* is a silent mutation, Y94Y, that does not alter the metabolizer phenotype, whereas *NAT2*7* results in an amino acid substitution, G286E, that leads to a significant decrease in the enzyme’s activity [12, 13]. The distribution of the NAT2 polymorphisms in the population studied were similar to other American populations in that one of the most frequent alleles was *NAT2*5*. It is established that the frequency of *NAT2*5* in European populations is ∼50%, in African populations is ∼33-42% and in Asian populations is

∼5% [14–17]. According to our results, the allele frequency of *NAT2*5* is ∼25%. The other two slow metabolizer alleles are *NAT2*6* and *NAT2*7*. The *NAT2*6* is common in all populations mentioned above with a frequency of ∼30%. Conversely, the frequency of *NAT2*7* is low in European populations (∼2%) and African populations (∼3-6%). In Asian populations, the frequency of *NAT2*7* is ∼10-12% [17]. Diverging from these reports, in our studied population the allele frequency of *NAT2*6* is ∼14% and of *NAT2*7* is ∼36%. As stated above, reduced NAT2 activity, which is observed in *NAT2*7* variants, can lead to adverse drug reactions due to increased accumulation of toxic metabolites. Additionally, our study revealed that the genotype frequency (predicted phenotype) of slow metabolizers is ∼47%. The relationship of NAT2 polymorphisms with INH-induced hepatotoxicity in TB patients among different populations were studied [14–21], but the previously published studies have demonstrated inconsistent results. Therefore, analysis of the slow genotypes should become part of the dosage regimen of INH in TB patients undergoing anti-TB treatment to prevent drug-induced liver injuries [18–21].

After NAT2 acetylates INH converting it to acetyl-INH, it can enter the CYP2E1 pathway, which couples with the glutathione-S-transferase (GST) metabolic pathway to facilitate the elimination of toxic metabolites [22–24]. Studies have shown that individuals with the *CYP2E1* wild-type allele (c1/c1 genotype) have a higher CYP2E1 activity that those with *CYP2E1*5B* allele (c1/c2 or c2/c2 genotype). Thus, these individuals can generate more hepatotoxins and therefore increase the risk of drug-induced liver injuries [23, 25, 26]. In our studied population, the allele frequency of the *CYP2E1*5B* is ∼ 79%, which increases the risk of hepatotoxicity specially in patients with a slow metabolizer phenotype for NAT2 [23, 27].

An important enzyme in the metabolism of RIF is AADAC catalyzing its deacetylation to 25-deacetyl-RIF [7, 28, 29]. Polymorphic variations affecting this enzyme’s activity can also result in the accumulation of toxic substances and variations in RIF plasma concentrations that can affect the efficacy of this drug. In this study, we analyzed the nonsynonymous SNP rs1803155 (AADAC*2 allele), which leads to a change in amino acid (V281I) in the coding region [30]. An allele frequency of ∼60% for *AADAC*2* has been reported in European American, African American, Japanese, and Korean populations. In our studied population, the allele frequency of *AADAC*2* is ∼99.9%. A limitation in this study is the number of SNPs analyzed in each gene, especially in AADAC. The analysis of additional genetic variations in AADAC can provide additional information in the metabolism of RIF. For example, the allele AADAC*3 (g.13651G>A/g.14008T>C), not analyzed in the current study, has shown a reduced metabolizing activity for RIF [30]. Additionally, studies have reported genetic polymorphisms in other RIF metabolizing enzymes, including carboxylesterase 1(CES1) and carboxylesterase 2 (CES2) [31], as well as in drug transporters and /or their transcriptional regulators, including SLCO1B1[32] and ABCB1 [33].

Countries have begun clinical trials focused on personalization of tuberculosis treatment to reduce the consequences for patients in treatment [21, 34]. In countries like Peru, where high rates of tuberculosis are recorded and therefore more people in treatment, the pharmacogenomic of individuals becomes a crucial tool for an optimum tuberculosis treatment. This review highlights the importance of having pharmacogenomic studies and having the identification of polymorphisms associated to the metabolism of the anti-tuberculosis drugs in our Peruvian population.

## Conclusion

In conclusion, our study showed the distribution of NAT2, CYP2E1 and AADAC genetic polymorphisms in a Peruvian population diagnosed with tuberculosis. This is a preliminary study to help understand the genetic basis of metabolizing polymorphisms in our population, and thus contribute to the use of this and future data in determining the safe INH and RIF dose in slow and fast metabolizers and thus minimizing adverse drug reactions. According to our results, ∼74% of the participants are carriers of slow metabolizer genotypes: *NAT2*5, NAT2*6* and *NAT2*7*, which increase the sensitivity of INH at low doses and increase the risk of drug-induced liver injuries. Additionally, ∼ 64% are homozygous for the wild-type CYP2E1*1A allele, which could increase the risk of hepatotoxicity. This high prevalence of slow metabolizers for Isoniazid in the Peruvian population should be further studied and considered to help individualize drug regimens, especially in countries with a great genetic diversity like Peru. However, 16% had a NAT2 fast metabolizer phenotype which could increase the risk of acquiring resistance to INH, thereby increasing the risk of multidrug-resistant (MDR) or treatment failure. The frequency of rs1803155 (AADAC*2 allele) was higher (99.9%) in Peruvians than in in European American, African American, Japanese, and Korean populations. These data will help the Peruvian National Tuberculosis Control Program develop new strategies for therapies.

## Notes

### Competing Interest Statement

The authors have declared no competing interest.

### Summary of Updates

this version clarifies the institutional affiliation of 2 authors

